# Head-direction coding in the hippocampal formation of birds

**DOI:** 10.1101/2020.08.31.274928

**Authors:** Elhanan Ben-Yishay, Ksenia Krivoruchko, Shaked Ron, Nachum Ulanovsky, Dori Derdikman, Yoram Gutfreund

## Abstract

Birds strongly rely on spatial memory and navigation. Therefore, it is of utmost interest to reveal how space is represented in the avian brain. Here we used tetrodes to record neurons from the hippocampal formation of Japanese quails – a ground-dwelling species – while the quails roamed a 1×1-meter arena (>2,100 neurons from 23 birds). Whereas spatially-modulated cells (place-cells, border-cells, etc.) were generally not encountered, the firing-rate of 12% of the neurons was unimodally and significantly modulated by the head-azimuth – i.e. these were head-direction cells (HD cells). Typically, HD cells were maximally active at one preferred-direction and minimally at the opposite null-direction, with preferred-directions spanning all 360°. The HD tuning was relatively broad (mean width ∼130°), independent of the animal’s position and speed, and was stable during the recording-session. Similarly to findings in rodents, the HD tuning usually rotated with the rotation of a salient visual cue in the arena. These findings support the existence of an allocentric head-direction representation in the quail hippocampal formation, and provide the first demonstration of head-direction cells in birds.

## Introduction

Since the seminal discovery of place cells in the rat hippocampus [1, 2], research on the mammalian hippocampal formation has been one of the most active fields in neuroscience. Half a century of extensive research resulted in a detailed characterization of hippocampal space processing in rodents, and advanced the development of new techniques and paradigms for neural recording in behaving animals, as well as new theories and ideas on the functional role of the hippocampus [3]. In addition to place cells, a whole range of other spatial cell types have been discovered in mammals, including head-direction cells [4], spatial-view cells [5], grid cells [6], border cells [7, 8], speed cells [9], and recently goal-direction cells [10]. The study of spatial processing in the hippocampus was not limited to rats but expanded to other mammals, including mice, bats, monkeys and humans [11-16]. The emerging notion is that the hippocampus and its related structures support spatial cognition and memory [17-19].

One important and relatively understudied question is whether the role of the hippocampal formation in spatial cognition is unique to mammals, or can we find its origins in other vertebrates? In this aspect, birds provide an interesting case study. Spatial navigation and foraging behaviors are common in avian species, and in some cases outperform those of mammals. Migratory and homing behaviors are stunning examples of bird navigational capabilities [20-24]. In adition, the extraordinary food caching and retrieval behaviors found in some song bird species demonstrate the impressive spatial-memory capacities of birds [25-27]. What might be the neural mechanisms that support such elaborate spatial cognition in birds? And how similar are they to the hippocampal space-processing system in mammals? Answers to these questions will constitute a breakthrough in our understanding of the evolutionary origin of space processing and, through comparative studies, may contribute to understanding the underlying mechanisms and to establishing general principles of navigation and spatial memory across vertebrates.

The avian hippocampal formation (HPF), a structure in the medial dorsal cortex of the avian brain (Fig. 1), is considered to be the homologue of the mammalian hippocampus, based on developmental, topographical, genetic, and functional lines of evidence[28-32]. Growing evidence suggest that it plays a role in spatial tasks [21, 24, 27, 28]. For example, experiments in HPF-lesioned pigeons consistently showed deficiencies in homing behaviors [33]. Similarly, lesion studies in zebra finches demonstrated that the HPF in these songbirds is involved in both learning and retention of spatial tasks [34]. Investigations on the patterns of activation of immediate early genes showed enhanced activation of the HPF during retrieval of cached food items [35] – a highly demanding spatial behavior – as well as during maze tasks [36]. Another striking indication for the involvement of avian HPF in spatial memory is found in the correlation between HPF volume and the importance of food caching for the natural behavior of the species [37]. Finally, neurons with place-fields were reported in the pigeon hippocampal formation, although these fields were mostly associated with rewarded locations [38-40]. These raise the hypothesis that the role of the mammalian hippocampus in spatial cognition, and the underlying neural spatial representation (place cells, grid cells, head-direction cells, etc.), have their roots in earlier vertebrate evolution. However, a clear allocentric representation of space, akin to that found in mammalian species, has not been reported yet in either the avian hippocampus, or in any other non-mammalian vertebrate.

**Figure 1.**
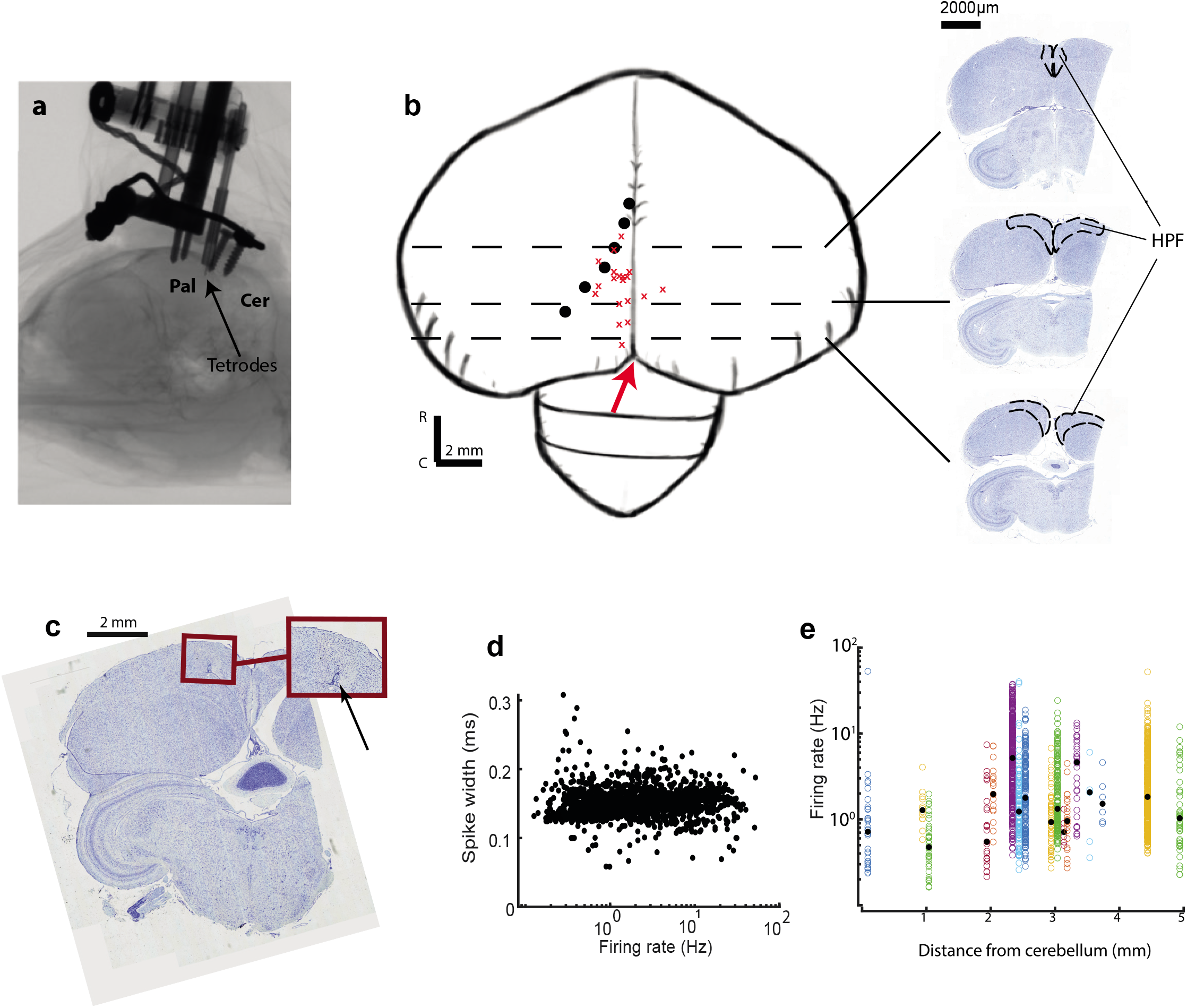
Recording locations and firing properties. **a**, Micro-CT scan showing an implanted microdrive on the quail’s skull. Arrow points to the tip of the tetrodes. Pal - pallium, Cer -cerebellum. **b**, A sketch of the quail’s brain. Red X’s mark the approximate recording locations from 19 quails. Black dots represent the putative lateral border of the hippocampal formation, which follows the lateral end of the lateral ventricle. Distances of black circles from the midline are approximated based on figures from the quail’s brain stereotaxic atlas [82]. Dashed lines indicate the approximate coronal section levels shown on the right. The dashed lines in the insets mark the hippocampal formation (HPF). The red arrow denotes the cerebellum tip. **c**, A Nissl stained coronal section showing an electrolytic lesion. Inset: magnified view of the lesioned region. The arrow points to the lesion mark. **d**, Spike width (full width at half maximum) for all the recorded single-units as a function of their firing rate. X-axis is logarithmic. **e**, Firing rates of single-units versus the recording location along the rostro-caudal axis (rostral distance from cerebellum tip). Colors designate different quails. Black circles designate the median firing rates. Y-axis is logarithmic.

In this study we report results of single unit recordings in the hippocampal formation of Japanese quails (*Coturnix japonica*). The Japanese quail is a ground-dwelling foraging bird, which has been extensively used as an animal model in developmental biology [41]. However, to our knowledge, single unit recordings in behaving quails have not been done before. The natural ground-foraging behavior of quails in relatively small areas is reminiscent of the rodents’ small-scale foraging behavior. We therefore asked if the resemblance of foraging behavior between quails and rodents implies resemblance in spatial representation. To this end, we used tetrode microdrives to perform electrophysiological recordings in freely behaving quails that explored a square arena, while we tracked their position and head-direction. Analysis of more than 2000 putative single-units in the HPF did not reveal clear spatially-modulated cells (e.g. place cells). However, we found that about 12% of the recorded cells showed a statistically significant (yet broad) head-direction response, which maintained stability over time, space, and across speeds. We thus provide the first evidence for a head-direction system in an avian species, shedding new light on the evolutionary and functional homology of the hippocampal system across taxa.

## Results

### No evidence for rodent-like place cells in the hippocampal formation of quails

We recorded and isolated 2,316 single units from 23 quails. In several of the quails, the signal-to-noise levels of the units were substantially reduced in the days following the surgery – hence the large variation in the number of cells collected from each quail – from 4 cells in quail 30 to 514 cells in quail 24 (Supplementary Table 1). Tetrodes were implanted in the left hemisphere in 21 out of the 23 quails and were stereotactically targeted to a zone within 5 mm rostrally from the cerebellar rostral tip, and within 2 mm laterally to the midline (Fig. 1a-b). 19 Penetration locations were reconstructed post-mortem by measuring the rostral and lateral distances of the penetration sites from the rostral tip of the cerebellum (Fig. 1b). In 4 quails the tetrode track was additionally reconstructed with an electrolytic lesion (Fig. 1c). In two of the birds penetration locations were reconstructed from microCT scans (Fig. 1a). Across the recorded population, firing rates and spike widths varied; but we could not detect systematic clustering of cell spike types (Fig. 1d) nor systematic variation of firing-rates along the rostro-caudal axis (Fig. 1e).

During the experiments, the quails sometimes explored the arena relatively evenly and sometimes in a restricted manner (Fig. 2a and Supplementary Fig. 1). In many cases, such as the example cell shown in Figure 2, the firing rate as a function of the quail’s position (rate map) displayed broad spatial firing across the arena (Fig. 2b). In this example cell, spatial information was not statistically significant (spatial information was smaller than the 99^th^ percentile of the shuffles). Only 19 cells (out of 938 cells that were recorded in behavioral sessions where the quail covered more than 50% of the arena) passed the criteria for place cells (spatial information larger than the 99^th^ percentile of the shuffles [p<0.01]). These neurons mostly did not display a clear single field of activity in their rate-maps (Supplementary Fig. 2). Therefore, we did not pursue further analysis of place modulations in this study.

**Figure 2.**
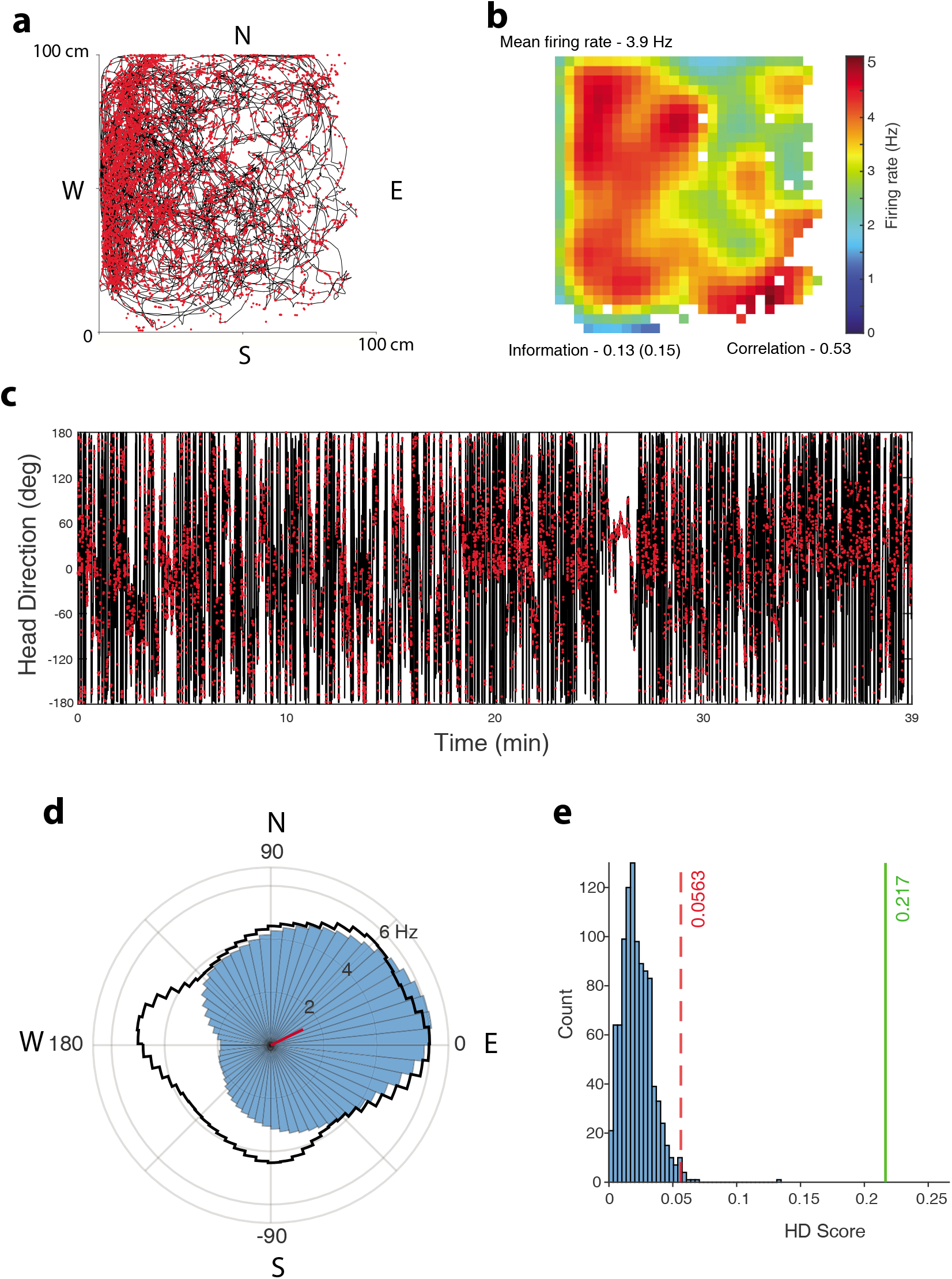
Data from an example neuron. **a**, The quail’s position in the arena during a 35-min recording session is marked by a black line. Red dots are the locations of spikes fired by the neuron. N - north side of the arena; S - south, E - east; W - west. **b**, Firing-rate map of the neuron. Indicated are the mean firing rate (top left), spatial information in bits/spike (bottom left), and map correlation between first and second half of the session (bottom right). White pixels are pixels which were not visited at least 200 ms during the session. **c**, Head-direction of the quail as a function of time (black line), with spikes superimposed (red dots). **d**, Polar plot showing the firing rate as a function of head direction (blue). The red bar indicates the length and direction of the Rayleigh vector. The black curve shows the relative time spent behaviorally in each head-direction bin. **e**, Rayleigh vector shuffling histogram. Red dashed line represents the Rayleigh vector length for the 99% percentile shuffle; green line shows the observed Rayleigh vector length for this example neuron.

### Head-direction cells in the hippocampal formation of quails

We next moved to analyzing the relationship between the firing rate and the head direction. In the example cell, firing rate was clearly modulated by the head direction (Fig. 2c), rising from 2 Hz at 180° azimuth to more than 6 Hz at 10° azimuth (Fig. 2d). The Rayleigh vector length (see Methods; termed thereafter ‘Rayleigh vector’) of the cell was 0.217; when compared to 1000 shuffled spike-trains, the observed Rayleigh vector was well above the 99^th^ percentile of the shuffled distribution (Fig. 2e) – and thus we categorized this example cell as having a significant HD response (p<0.01). The preferred direction of the response (computed as the direction of the Rayleigh vector) pointed roughly North-East (red bar in Fig. 2d).

Out of the recorded population, 260 cells (∼12%) passed the shuffling statistical test as head direction cells (with a chance level of 1%; Supplementary Table 1). To confirm that the number of HD cells is not biased by the parameters used to define a single-unit (isolation distance and L-ratio criteria; see Methods), we analyzed the percentage of HD cells at 8 different combinations of criteria (Supplementary Table 2). The value of roughly 12% HD cells was maintained regardless of the combination of criteria used.

### Stability and distribution of preferred directions

In general, the HD modulation tended to be relatively broad (Fig. 3a). The mean tuning-width (width at half height, measured halfway between maximum and minimum of the tuning) for the population of significant HD cells was 130.4° ± 54° (Fig. 3b; mean ± s.d.). Moreover, the modulation depth (normalized difference between the highest and lowest firing rate) varied between 0.4 and 1, with a mean of 0.66 (Fig. 3c). To compare these mean values with the corresponding values of head direction cells in rodents, we performed the same analyses on datasets of previously recorded neurons from the medial entorhinal cortex (MEC), parasubiculum (PaS) and the dorsal presubiculum (dPrS) of rats, areas that are well known for their robust representation of head direction (data from [42] and [43]). Out of the 2404 cells from rats that we analyzed, 1046 were significant HD cells, comprising 44% (compared to 12% in the quails’ HPF). The distribution of widths at half height, the standard deviations of the rate curves, and the Rayleigh vectors of HD cells – all pointed to substantially narrower tuning of rat HD cells as compared to quail HD cells (Supplementary Fig. 3). Moreover, the mean modulation depth of the HD cells in rats was substantially larger compared to the HD cells in quails (0.85 compared with 0.64 in quails; Supplementary Fig. 3).

**Figure 3.**
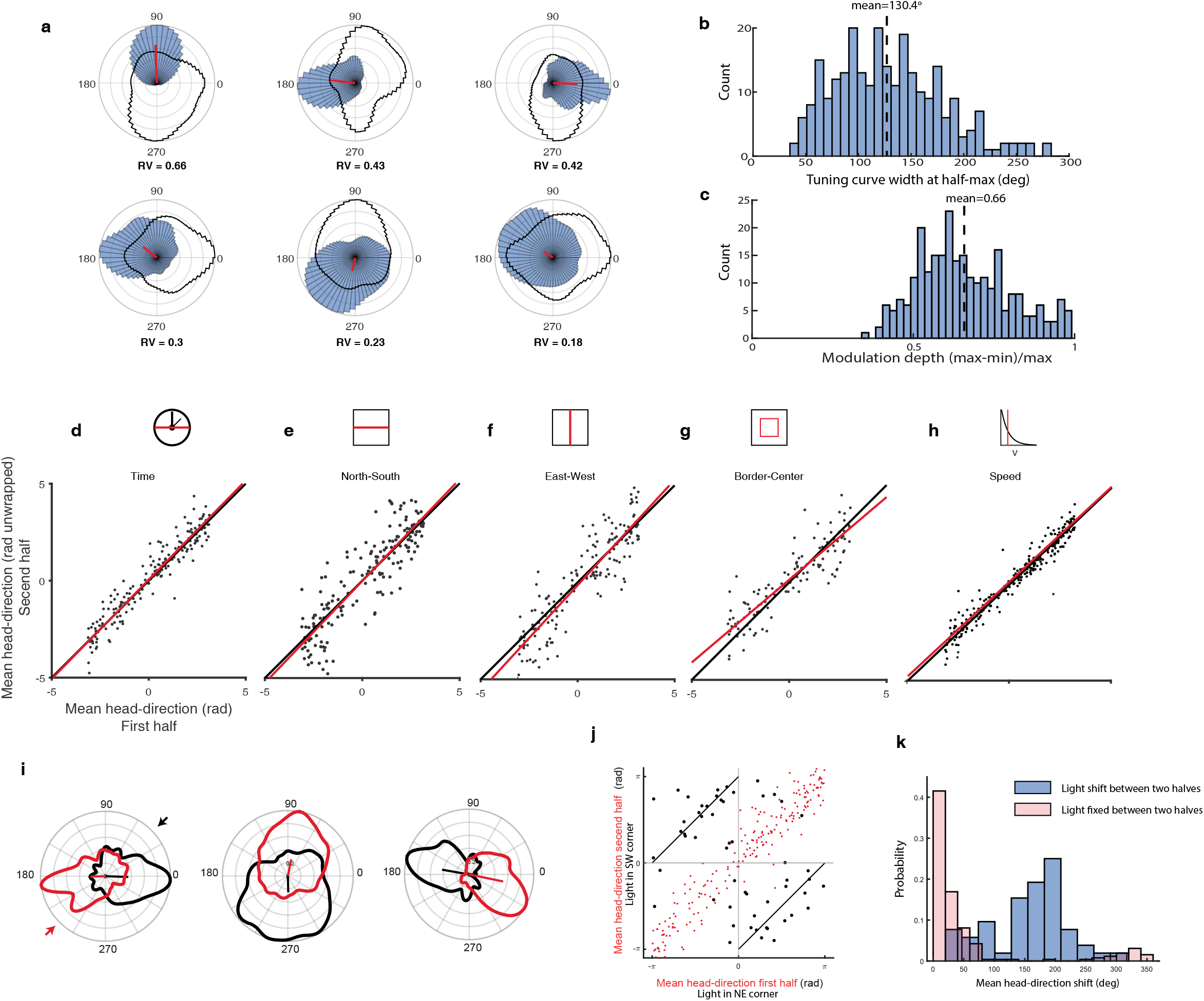
Head-direction cells characteristics, stability, and response to cue-rotation. **a**, Examples of six cells significantly modulated by head-direction, organized in descending order by their Rayleigh vector (RV; its value is indicated below each neuron). Blue histograms represent the polar firing-rate curve. Black line depicts the time spent in each direction (behavioral curve). Curves are normalized to their maximum. **b**, Distribution of the tuning curve widths (width at half maximum) of the head direction firing-rate curves. The dashed line represents the population mean. **c**, Distribution of the modulation depths (maximal firing-rate of the tuning curve minus the minimal firing rate, normalized by the maximal firing rate). The dashed line represents the mean. **d-h**, Population scatter plots for HD cells, comparing the preferred head-directions in the first half versus the second half of the session (d), The north versus south parts of the arena (e), east versus west parts of the arena (f), near-border versus center of the arena (g), and fast speeds versus slow speeds (h). Black lines are the identity-lines (equal preferred-directions) and red lines show the linear regressions. **i**, Examples of head-direction tuning curves of three cells before and after 180°cue rotation (light source switch). Red curves were measured when the light source was in the South-West corner of the arena and black curves when the light source was in the North-East corner (the inset arrows in the left panel designate the directions of the light sources in the arena. Straight lines originating from the center designate the direction and length of the Rayleigh vectors. **j**, Preferred head-directions in the second half of the session are plotted versus the preferred head-directions in the first half of the session. The black dots show the results when the light source was shifted at mid-session by 180°. For comparison, superimposed in red are the dots from panel d to show the result without a light shift. The black diagonal lines indicate the predicted 180o shift of the preferred direction. **k**, Distribution of the differences between preferred head directions measured in the two halves of the session, when the light was shifted by 180⊠ (blue) compared to no shift (red).

Despite their relatively broad shape, the HD tuning curves in the quails had high directional stability: (i) The preferred directions in the first and second halves of the session were significantly correlated (Fig. 3d; Pearson correlation: r=0.95, p<0.0001, n=186; the numbers of neurons in this and following comparisons is lower than n=260 because we required sufficient coverage in both halves: see Methods). (ii) The preferred directions computed from the north versus south halves of the arena were significantly correlated (Fig. 3e; Pearson correlation: r=0.88, p<0.0001, n=175), and likewise for preferred directions from the west versus east halves (Fig. 3f; Pearson correlation: r=0.90, p<0.0001, n=169). (iii) Because some of the quails mostly explored the borders of the arena (see examples in Supplementary Fig. 1), we also analyzed the preferred head directions when the quails were near the border versus when they were in the center of the arena; these were also found to be highly significantly correlated (Fig. 3g; Pearson correlation: r=0.88, p<0.0001, n=122). (iv) Finally, the preferred directions during fast movement of the quail (>5 cm/sec) were significantly correlated with the preferred directions during slow movements (<5 cm/sec) (Fig. 3h; Pearson correlation: r=0.97, p<0.0001, n=260). In all these tests the regression line was very close to the 45° line (Fig. 3d-h: compare red and black lines). Our results indicate a stable head-direction representation over time and space, as well as over the speed of the quails. The significant correlation between the preferred directions in different parts of the arena argue against the possibility that the head-direction tuning is an outcome of spatial view cells [5].

In rodents, HD cells commonly rotate their preferred direction following rotation of salient cues in the arena [4, 44]. We therefore positioned, in some of the experiments, two LED light-sources in opposite sides of the arena, have lit only one of them, and then switched the side of the illuminating source at mid-session. 52 significant HD cells were analyzed in this experiment. Following the switch, some of the neurons demonstrated clear 180° rotation of their tuning curve (Fig. 3I). At the population level, the distribution of the preferred-direction differences between light on one side and light on the 180°-opposite side was centered near 180° (Fig. 3J and 3K; mean angle ± circular standard deviation = 162±64°), and significantly deviated from the distribution of the mean directional shift between the two halves of the session without any cue-changes (Fig. 3J and 3K; Watson-Williams mean angle test, p<0.0001).

The preferred directions of the population covered almost uniformly the entire range of 360°)Fig. 4a). This uniformity was seen when pooling data from all quails (Fig. 4a), as well as in most of the individual quails (Supplementary Fig. 4). Significant HD cells, spanning all possible directions, have been recorded at all the recording-depths in the HPF and at all anatomical locations within HPF (Fig. 4b-c). HD cells were found at nearly all depths from the brain-surface, and all distances rostral to the cerebellum tip (Fig. 4d-e). Thus, our results do not indicate a clear tendency for anatomical clustering of HD cells or anatomical clustering of best directions. We conclude that significant and stable HD cells are found throughout the HPF of quails.

**Figure 4.**
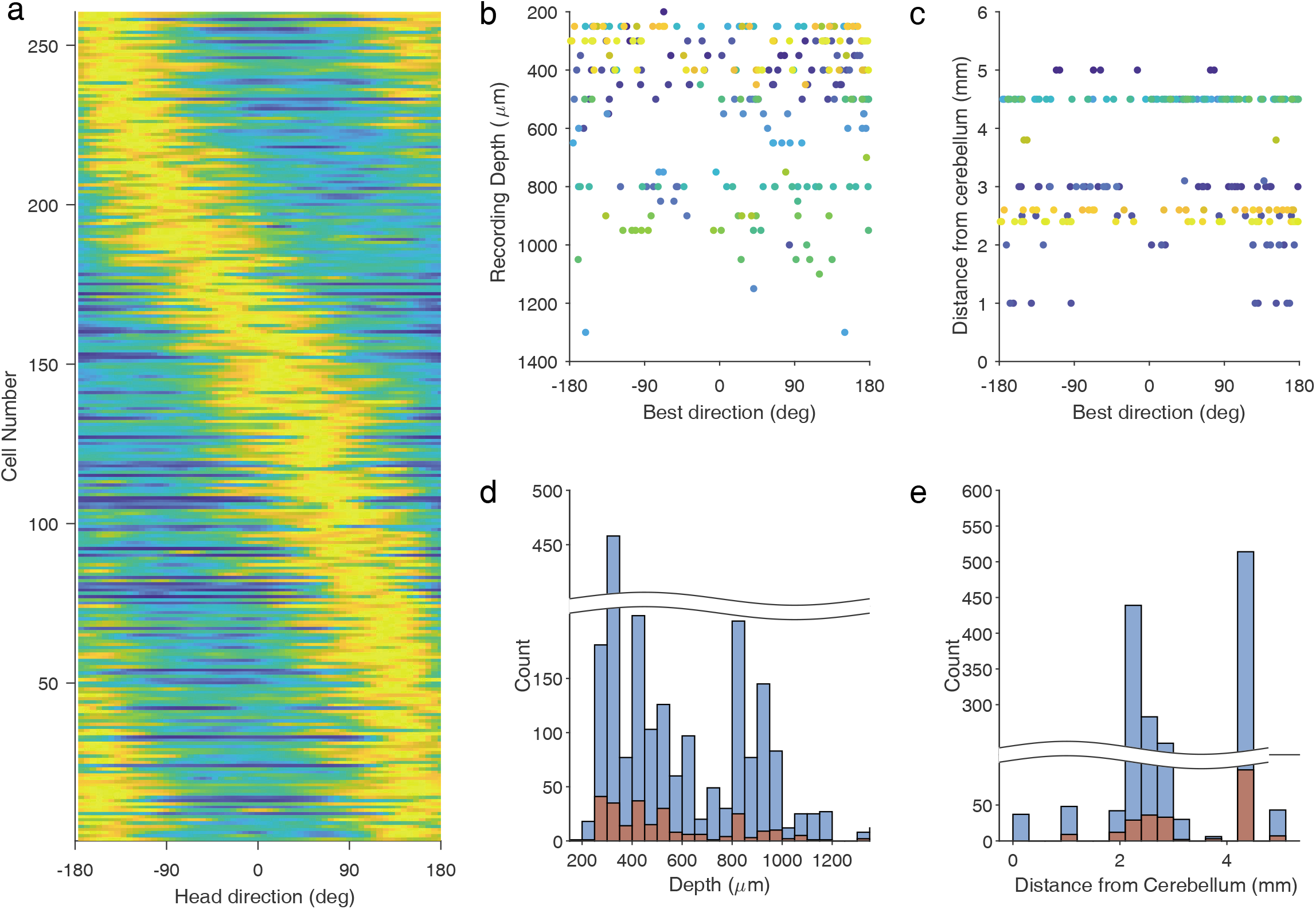
Distributions of preferred directions across the population of head-direction cells. **a**, Normalized HD tuning-curves for all significant head-direction cells, ordered according to the preferred direction. **b**, Preferred directions of HD cells as a function of estimated recording depth. Colors designate different quails. **c**, Preferred directions as a function of recording distance from the cerebellum along the rostro-caudal axis. Colors indicate different quails. **d**, Histograms of total number of single units recorded (blue) and number of HD cells (orange) as a function of recording depth. **e**, Same as d, but plotted as a function of recording distance from the cerebellum.

## Discussion

To our knowledge, this is the first time that a stable and abundant population of head-direction cells is reported in the hippocampal formation of an avian species. Neurons that represent head direction in allocentric coordinates during real-world movements have been reported in mammals [4, 45-47], and very recently in fish [48] – as well as in head-fixed insects in virtual reality [49, 50]. These cells are thought to underlie the directional sense of animals, which is vitally important for goal directed navigation and other spatially coordinated behaviors [51]. Because many species of birds display remarkable cognitive capabilities that are related to spatial behaviors and spatial memory [20-24, 26], an elaborated head direction system is expected in avian brains. However, up until now such a system has not been discovered. A characteristic of mammalian systems representing head direction is that they exhibit a uniform and stable representation of directions [51, 52]. Moreover, direction sensitivity is maintained independent of the speed of the animal [4]. Here we report a population of neurons that are modulated by the head direction of the quail: these neurons uniformly represent all directions, and their preferred direction is stable across time, space, and movement speeds. We therefore posit that these cells are part of a head-direction system in quails, perhaps homologous to that of mammals.

Networks of head-direction cells in mammals integrate angular velocity information coming from the vestibular system with environmental cues from the visual system [53]. This integration is believed to take place in the dorsal presubiculum [54], anterodorsal thalamus, and lateral mammillary nucleus [55]. The head-direction information is conveyed from the dorsal presubiculum to the entorhinal cortex, where it interacts with other spatial information coming from the hippocampus [56, 57]. The sub-division homology between the avian HPF and the mammalian HPF, is not well established [28]. Atoji and Wild (2004, 2006) provided a histochemical division of the pigeon HPF to dorsal lateral (DL), dorsal medial (DM) and a medial V shape complex [30, 58]. The homology of this subdivision with the mammalian HPF is debated, with the controversy focused mostly around the homologies of the V-shaped complex and the dorsal medial parts with the mammalian Dentate Gyrus and Ammon’s horn [59-63]. However, there seems to be an agreement regarding the avian DL being related to the mammalian subiculum and entorhinal cortex [28, 30, 64-66]. Many of the HD cells that we recorded were in the more accessible dorsal lateral (DL) and dorsal medial (DM) parts of the HPF (Fig. 1b and Fig. 4b) – thus, our findings possibly represent an avian head-direction system homologous to the subiculum/entorhinal HD system of mammals. Notably, the DL and DM portions of the HPF receive connections from visual Wulst [67] as well as connections from the dorsal thalamus [68], which may provide the anatomical substrates for integration of visual and self-motion cues. Yet, obvious differences exist between our findings and analogous reports in rodents: (1) Most notably, many of the HD cells in the entorhinal cortex and subiculum of mammals fire close to zero spikes (< 0.5 Hz) in the null directions [69, 70], whereas in our population of HD cells the neurons had a non-zero baseline firing at all head directions, with significantly higher rates in the preferred direction as compared to the null directions. (2) Moreover, in the subiculum and medial entorhinal cortex of rodents many of the neurons are narrowly tuned (<45° width at half peak) [69]. Such narrowly tuned neurons were a small fraction of our population of HD cells – while most of the quail’s HD cells were more broadly tuned. (3) In addition to the above-mentioned differences, the abundance of HD cells in the avian HPF was lower than in the mammalian subiculum and layers III to VI of medial entorhinal cortex, where more than 40% of the neurons are significant HD cells [43, 70]– compared to about 12% of the recorded cells in the quail. These differences between findings in quails and rodents are not a reflection of different metrics used in different papers, because we repeated the analysis on raw-data of population of neurons recorded from rats in the Pas, dPrS and MEC – using the exact same analysis-pipeline – and have confirmed the substantial differences in modulation depth, tuning broadness and abundance of HD cells (Supplementary Fig. 3). The relatively low abundance of HD cells in our findings may reflect the non-focal sampling of neurons in our study: Perhaps a higher fraction of HD cells would be found in specific sub regions of the HPF. However, our results showed no evidence for anatomical clustering of HD cells in the quail’s HPF (Fig. 4d-e).

Functional hemispheric lateralization is common in birds with laterally-placed eyes [71]. Recordings in the pigeon HPF demonstrated a significant difference in the behavior of neurons between the two hemispheres: neurons in the left HPF tended to code spatial parameters more reliably, while neurons in the right HPF tended to code goal locations [40]. Lateralization of HPF spatial coding is also supported by a lesion study in pigeons showing that an intact left but not right HPF is required for using local landmarks to guide food search [72]. Following these studies, we performed most of our recordings in the left hemisphere. However, we did record in two quails from the right hemisphere (quails # 10 and 23; Supplementary table 1) and found 11 HD cells out of 123 cells (9%). Thus, we have no indication for lateralized organization of HD cells in the quail. However, further sampling is required for a thorough comparison between hemispheres.

Another striking difference between neural responses in rodents and the results reported here is the apparent lack of clear place cells, grid cells and border cells. A small number of neurons showed spatial modulation of their firing rates, but these were mostly broadly tuned (Supplementary Fig. 2). These results are consistent with results reported by Bingman and colleagues [38-40], who studied the HPF of pigeons in radial mazes and open arenas. These previous studies found cells that were significantly spatially modulated, but in many cases the responses were consistent with multiple broad fields and were related mostly to rewarded locations. Particularly, in pigeons foraging in open arenas without stable goals (comparable to our experiment), spatially modulated signals were scarce [39]. A preliminary electrophysiological survey of Zebra Finch HPF also identified scarce place-cells with relatively broad spatial tuning [73]. On the other hand, behavioral experiments in quails [74], pigeons [75] and zebra finches [76] strongly support well-developed spatial cognitive abilities in these bird species – such as novel shortcuts, model-based spatial learning, navigation, and spatial memory. It is possible that the neural substrate of spatial perception of birds is not localized in the HPF, or alternatively, it is localized in a specific sub-region of the HPF which has not been explored yet. However, lesion studies in quails, pigeons and zebra finches [34, 77, 78], as well as gene expression analysis [79, 80], provide strong evidence that the HPF is critically important for spatial cognition. A different possibility is that the HPF of both mammals and birds holds the neural basis of spatial cognition but evolved differently: rather than having a large population of highly-specific and sparsely encoding cells – as seen in mammals – birds might possess a population-coding scheme which is carried by a population of neurons with broad tuning and non-uniform fields. This possibility is consistent with our findings of head direction tuning which was broadly tuned.

A preliminary study in food caching birds that are specialized in spatial memory (tufted titmouses) revealed place cells in the rostral HPF [73], in the same areas that have been explored in other birds, including here in quails – but without observing place cells. Taken together, our and previous findings suggest the notion that the HPF in birds generally supports spatial cognition – but that the detailed coding scheme is species-specific and evolved differently according to the ecological needs of the species. Previous studies of hippocampus activity in freely behaving pigeons did not track the head-direction[39] and therefore it is not known if HD cells can be found in other species of birds. However, in the above-mentioned preliminary study of a food caching species, head direction was tracked, and HD cells were observed in addition to place cells [73]. Our study is thus unique to report HD cells in an avian species lacking place-cells. This is an interesting observation and taken together with observations of HD cells in a variety of species – ranging from insects, fish and birds – it suggests that HD cells, and not place cells, were conserved through evolution. These intriguing inter-species differences call for further comparative investigations of spatial cell types across vertebrates and linking it to the animals’ behavioral repertoire and evolutionary history: A merging of neuroscience, evolution, and behavioral ecology.

## Methods

### Animals

Japanese quails (*Cotrunix japonica)* of both sexes were used in this study. The quails were hatched and raised in our in-house breeding colony, housed in 1×1 m cages and maintained on a 12/12hr light/dark cycle. Food and water were provided *ad libitum*. All procedures were in accordance with the ethical guidelines and were approved by the Technion Institutional Animal Care and Use Committee. During recording sessions, no painful procedures were carried out.

### Surgery

Adult Japanese quails (females and males; weight: 150 – 250 g, age: 4 – 12 months old) were prepared for repeated electrophysiological recordings with a single surgical procedure: the birds were anaesthetized with 3% isoflurane in a 4:5 mixture of nitrous oxide and Oxygen. Quails were then positioned in a stereotaxic frame (Kopf, small animals instrument Model 963) using Kopf rat ear bars. Head angle was controlled with a biting bar that was positioned 45° below the inter-aural axes (resembling the standard position in the pigeon and the quail atlases [81,82). At this head position, in our stereotaxic setup, the cerebellum rostral tip was found at coordinate 0 (inter-aural line) ±0.5 mm rostro-caudal. During surgery, animal temperature was maintained using a closed-circuit heating pad. Lidocaine (Lidocaine HCl 2% and Epinephrine) was injected locally at the incision site. The skull was exposed and cleaned. Two to four skull screws were inserted at the caudal part of the skull and one ground screw was inserted at the right frontal part of the skull. A 2 mm craniotomy was performed, in various positions ranging up to 4 mm rostral from the inter-aural line, and a small nick in the dura was made at the center of the craniotomy with a surgical needle. A custom-made Microdrive (as described in {Weiss, 2017 #1587]) containing four tetrodes (made of 17.8-μm platinum-iridium wire, California Fine Wire) was carefully lowered until the tetrodes smoothly entered the brain tissue. Antibiotic ointment (Chloramfenicol 5%) was applied to the brain surface, followed by a thin silicon coat (Kwik-Sil). The drive was then connected to the ground screw with a silver-wire and attached to the skull with adhesive cement (C&B-metabond) and light cured dental resin (Spident Inc. EsFlow). Loxicom (5mg/ml) was injected intramuscularly and the quails were positioned in a heated chamber to recover overnight.

### Electrophysiological recordings

Electrophysiological recordings were conducted in a 1 x 1 m open-field arena. The birds were released to roam spontaneously in the arena. Occasionally, food items were scattered in the arena to encourage movement. A tethered 16-channel headstage (Intan RHD2132) was attached to the drive and connected through a commutator (Saturn) to a data acquisition system (Neuralynx Digital SX). In some of the experiments the quails were untethered, in which case a 16-channel neural logger (Deuteron Technologies, MouseLog-16) was used for on-board acquisition and data storing. In both cases, animals were filmed with a CCTV camera with a frame-rate of 25Hz, and their position and head orientation was tracked by video-tracking two LEDs (green and red) mounted on both sides of the headstage or the neural-logger. Recording sessions were performed daily and lasted 10–45 minutes. Tetrodes were advanced by 50 μm per day. If no spikes were detected, electrodes were continuously lowered until spikes were observed. After reaching a depth of more than 1000 μm, in some of the experiments, electrodes were retracted to regions of previously observed spiking activity and additional experiments were conducted. The thickness of the quail HPF layer varies from ∼800 μm on the lateral side to ∼2000 μm on the medial side[82]. Therefore recording depth was restricted to within 1200 μm below the brain surface, with more than 80% of the neurons collected in depths between 200 to 800 μm (Fig. 4b). Thus, the bulk of the recorded population of neurons was collected within the HPF. Yet, we cannot rule out the possibility that a small subset of the neurons from deep sites were recorded below the lateral ventricle, or that neurons in the most lateral penetrations were from bordering areas adjacent to the HPF.

Electrical recording was sampled at 30 kHz using the Cheetah 6.0 software (Neuralynx), which records both 16-channel continuous raw data and spike waveforms. Raw signals were pre-amplified, bandpass filtered to 300 – 3000 Hz and thresholded (30 – 60µV) to obtain on-line spiking activity (see Supplementary Fig. 5 for examples of filtered traces). In untethered recordings, signals from the 16 channels of the neural-logger were amplified (×200), filtered (300 – 7000 Hz) and sampled continuously at 29.3 kHz or 31.25 kHz per channel, and stored on-board. For synchronization between the neural-logger data recording and the Neuralynx video recording, a neural-logger control box sent TTL pulses to both the neural-logger and Neuralynx system at random intervals (10 ± 5 seconds). The timestamps were synchronized by utilizing polynomial curve fitting between the pulses recorded on both systems.

### Data processing and spike sorting

During subsequent processing (off-line), the electrical recordings were filtered between 600 – 6,000 Hz, and an adaptive voltage threshold (Thr) was used for spike detection, as describe by [83]:

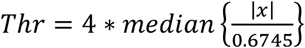

Where *x* is the filtered recorded signal, the division by 0.6745 accounts for the relationship between the median of the noise and the standard deviation of a normal distribution. Voltage threshold was calculated over 1-minute recording windows. Electrical artifacts were detected and removed from all channels based on absolute voltage. Following artifact removal, a spike detection algorithm was used for each tetrode: whenever one channel crossed the threshold value, a 1-ms wide segment was saved from all channels on the tetrode, where the peak was centered over the 8^th^ sample (for a total of 32 samples). Coincidence detection algorithm was used to further identify and remove movement artifacts; large-amplitude events occurred within all channels in each tetrode or between tetrodes were removed from analysis (Supplementary Fig. 5b). Additionally, all detected spikes were compared to a pre-existing spike-shape database (*template matching*) by correlating different segments of the spike shape to each template from the database. A spike was included for further analysis if its Pearson-correlation coefficient with any of the spike templates was larger than 0.8.

Manual spike sorting was performed offline using SpikeSort3D software (Neuralynx) and consisted of plotting the spikes in 3D parameter space, finding the features which give the best cluster separation and performing manual cluster cutting. Clustering quality was assessed based on two measures: isolation distance and L_ratio_ [84]. These parameters estimate how well-separated is the cluster of spikes from the other spikes and noise recorded on the same tetrode. A well-separated cluster has a large isolation distance and a small L_ratio_. A cluster was considered a single unit if it had an L_ratio_ smaller than 0.2 and an isolation distance larger than 15. When the number of points outside the isolated cluster is smaller than half of the total points in the tetrode it is not possible to obtain the isolation distance value (NaN isolation distance). Clusters with NaN isolation distance were included in the analysis (see Supplementary Table 2). Additionally, clusters were categorized as multi- or single-units based on their inter-spike interval histogram: a cluster was categorized as single-unit if less than 5% of its spikes occurred less than 2 ms after the previous spike.

### Histology

In a few of the quails, an electrical lesion was performed by injecting a positive current through one of the tetrodes (+5μA for 20 sec). A week later, the quail was deeply anesthetized and perfused with phosphate buffer solution (PBS) followed by 4% paraformaldehyde. The brain was removed and stored in 4% paraformaldehyde for 2–3 days at 4°C, then transferred to PBS. Following fixation, the quail brains were dehydrated in 70%, 80%, 95% and 100% ethanol, cleared in Xylene and embedded in paraffin wax. The paraffin-embedded brains were coronal sectioned at 5 µm using a microtome (RM 2265 Leica). Sections collected at 40 μm intervals were mounted on superfrost glass slides and dried in an oven at 37°C for 24 hours. After drying the sections, they were deparaffinized in xylene, rehydrated in a diluted ethanol series, and stained with 0.1% Cresyl violet solution (Nissl stain). The sections were then dehydrated, cleared and cover-slipped with DPX mounting medium (Merck).

### Data analysis

Single units were included for further analysis only if at least 300 spikes occurred during the session, and the recording session lasted more than 10 minutes.

Spatial firing-rate maps were computed by partitioning the arena into 3×3 cm bins. The number of spikes in each bin was divided by the time the bird spent in that bin. Bins which the quail visited for less than 200 ms were discarded and were colored white in the firing-rate map. Experiments in which the quail visited less than 50% of the arena were discarded from the spatial rate-map analysis. Spatial modulation was assessed based on two common measures: spatial information and spatial correlation [85]. Spatial information index was calculated as:

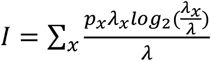

Where *λ* is the mean firing rate of the cell, *λ*_*x*_ is the mean firing rate in bin *x* (x includes only bins that the quail visited), *p*_*x*_ is the probability of occupying bin *x*. Spatial correlation was the 2-dimensional correlation coefficient between the rate map of the first and second halves of the session – an index of stability. For statistical evaluation of the spatial modulation, a shuffling procedure was applied. The entire spike train was rigidly and circularly shifted by a random interval (intervals smaller than ±20 s were not used). This was repeated 1000 times for each neuron, and for each of the shuffles the spatial information index was calculated. The experimentally observed index for the actual neuron was considered statistically significant with a P value larger than 0.01 if it surpassed 99% of the shuffle indices. For display purposes, the rate maps were smoothed using a 2D Gaussian kernel (σ = 1 bins), but all the computations and indexes were calculated without smoothing.

Head-direction was computed as the perpendicular orientation from the line connecting the positions of the two LEDs (we used the red and green colors of the LEDs to unambiguously determine the correct direction). Head direction data were binned at 6° bins. The number of spikes in each bin was divided by the time the animal spent at that direction. The histogram was smoothed using a hamming window (n = 10 points) and displayed in polar plot view.

To assess the directionality of the tuning curve, the mean vector length of the circular distribution (Rayleigh vector) was calculated:

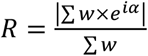

Where *w* is the firing-rate per bin and *α* is the bin direction. The preferred (mean) direction of the tuning curve was estimated by the direction of the Rayleigh vector:

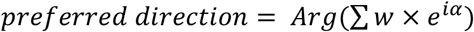

The statistical significance of the directionality was assessed by applying a spike-shuffling procedure as described above. Shuffling was repeated 1000 times, and for each shuffle the Rayleigh vector (RV) length was calculated. A neuron was considered head-direction modulated if the RV score was larger than 99% of the shuffled scores. A shuffling procedure was used only for tuning curves with a RV score higher than 0.10.

The stability of the HD preferred direction was assessed by dividing the time of the experiment into two halves (Fig. 3d); by dividing the arena into two regions: south half versus north half, east half versus west half, and center region versus border region (Fig. 3e-g); and by dividing the quail’s speed to two speeds: below 5 cm/sec and above 5 cm/sec (Fig. 3h). Preferred HD was calculated separately for each part. This analysis was performed only for the sessions where the head direction coverage was larger than 50% in both regions. Head direction coverage was calculated by dividing the time spent at each region by 60 (the number of 6° HD bins) and thus obtaining the expected time at each direction-bin if the behavioral head directions were uniformly distributed. The percent coverage was then defined as the percentage of directional bins in which the quail spent more than 75% of the expected uniform time (see Supplementary Fig. 1 for examples).

## Acknowledgments

We thank Dr. Yael Zahar for assistance and support, and May-Britt Moser, Edvard I. Moser and Charlotte N. Boccara for usage of rat data in Supplementary Fig. 3. This work was supported by research grants from the Rappaport Institute for Biomedical Research, the Adelis Foundation and the Israel Science Foundation. Yoram Gutfreund also acknowledges the generous support of the Edward S. Mueller Eye Research Fund.

## Author contributions

E.B., K.K. and S.R. performed the experiments, wrote code, analyzed data and produced graphs and figures. N.U., D.D. and Y.G. planed the experiments and designed the experimental setups. All authors contributed to the writing of the manuscript.

## Notes

### Competing Interest Statement

The authors have declared no competing interest.

### Summary of Updates

Additional experiments were performed to test the stability of HD cells, by presenting and rotating salient cues within the arena and testing for the corresponding rotations of the cell's tuning curve. Additionally, the parameters of the Quail's HD cells were compared to similar cells found in Rats.

