## Supplementary Material for "Head-direction coding in the hippocampal formation of birds"

#### Supplementary Tables

**Table S1. Summary of recorded cells for individual quails.** Locations of penetrations relative to the cerebellum were measured post mortem.

| | | Location of penetration | | Modulation Type | | | | |
|---|---|---|---|---|---|---|---|---|
| | Quail# | From Cerebellum (mm) | Lateral from midline (mm; right positive) | Head direction cells | Place cells | Speed cells | Border cells | Total cells |
| **1** | 1 | 0.15 | -0.4 | 0 | 0 | 0 | 2 | 32 |
| **2** | 2 | 3 | -0.5 | 0 | 0 | 0 | 0 | 6 |
| **3** | 3 | 5 | -0.68 | 7 | 0 | 0 | 0 | 43 |
| **4** | 4 | 3 | -0.5 | 5 | 0 | 0 | 0 | 22 |
| **5** | 5* | 3 | -0.1 | 1 | 0 | 0 | 0 | 5 |
| **6** | 6 | 3 | -1 | 19 | 1 | 0 | 1 | 54 |
| **7** | 7 | 2 | -0.3 | 3 | 1 | 0 | 1 | 25 |
| **8** | 9 | | | 2 | 0 | 0 | 3 | 27 |
| **9** | 11 | 1 | -0.3 | 5 | 0 | 0 | 1 | 36 |
| **10** | 15 | 2.5 | 1.5 | 10 | 2 | 0 | 1 | 115 |
| **11** | 17 | 1 | -0.5 | 4 | 0 | 0 | 0 | 12 |
| **12** | 18 | 2 | -0.5 | 9 | 0 | 0 | 0 | 17 |
| **13** | 20 | 3.1 | -1 | 2 | 0 | 0 | 0 | 30 |
| **14** | 22* | 3 | -0.6 | 8 | 0 | 2 | 2 | 164 |
| **15** | 24* | 4.5 | -1 | 99 | 1 | 1 | 7 | 514 |
| **16** | 27 | | | 18 | 8 | 0 | 4 | 305 |
| **17** | 30 | | | 2 | 0 | 0 | 0 | 4 |
| **18** | 31 | 3.8 | -1.6 | 3 | 0 | 0 | 0 | 6 |
| **19** | 32 | | | 8 | 0 | 1 | 2 | 94 |
| **20** | 33* | 2.6 | -1.5 | 26 | 16 | 0 | 4 | 179 |
| **21** | 36 | 2.4 | -1.6 | 29 | 3 | 9 | 18 | 442 |
| **Total** | | | | **260** | **32** | **13** | **46** | **2127** |
| **22** | 66** | 3.0 | -0.3 | 33 | 14 | - | - | 181 |
| **23** | 70** | 2.5 | 0.4 | 1 | 0 | - | - | 8 |

* - Location verified from electrode traces in Nissl stained slices

** - Location verified with a micro CT scan. Data not included in the cell summary (descriptive statistics in Fig. 3 a-h and Supplementary Fig 3)

**Table S2. Percent head direction cells based on different spike isolation criteria.** The percentage of HD cells has been calculated using eight different combinations of criteria for single unit isolation: Upper limit of L-ratio, Lower limit of Isolation Distance and whether to include clusters with NaN isolation distance values (when the number of points outside the isolated cluster is smaller than half of the total points in the tetrode an isolation distance value cannot be computed).

| L-ratio | Isolation distance | NaN values | Cells total | HD Cells | HD cells percentage [%] |
|---|---|---|---|---|---|
| 0.2 | 15 | included | 2144 | 270 | 12.6% |
| 0.2 | 30 | included | 1354 | 168 | 12.4% |
| 0.1 | 15 | included | 1674 | 201 | 12.0% |
| 0.1 | 30 | included | 1287 | 157 | 12.2% |
| 0.2 | 15 | not included | 1761 | 217 | 12.3% |
| 0.2 | 30 | not included | 971 | 115 | 11.8% |
| 0.1 | 15 | not included | 1305 | 151 | 11.6% |
| 0.1 | 30 | not included | 918 | 107 | 11.7% |

Supplementary figure 1

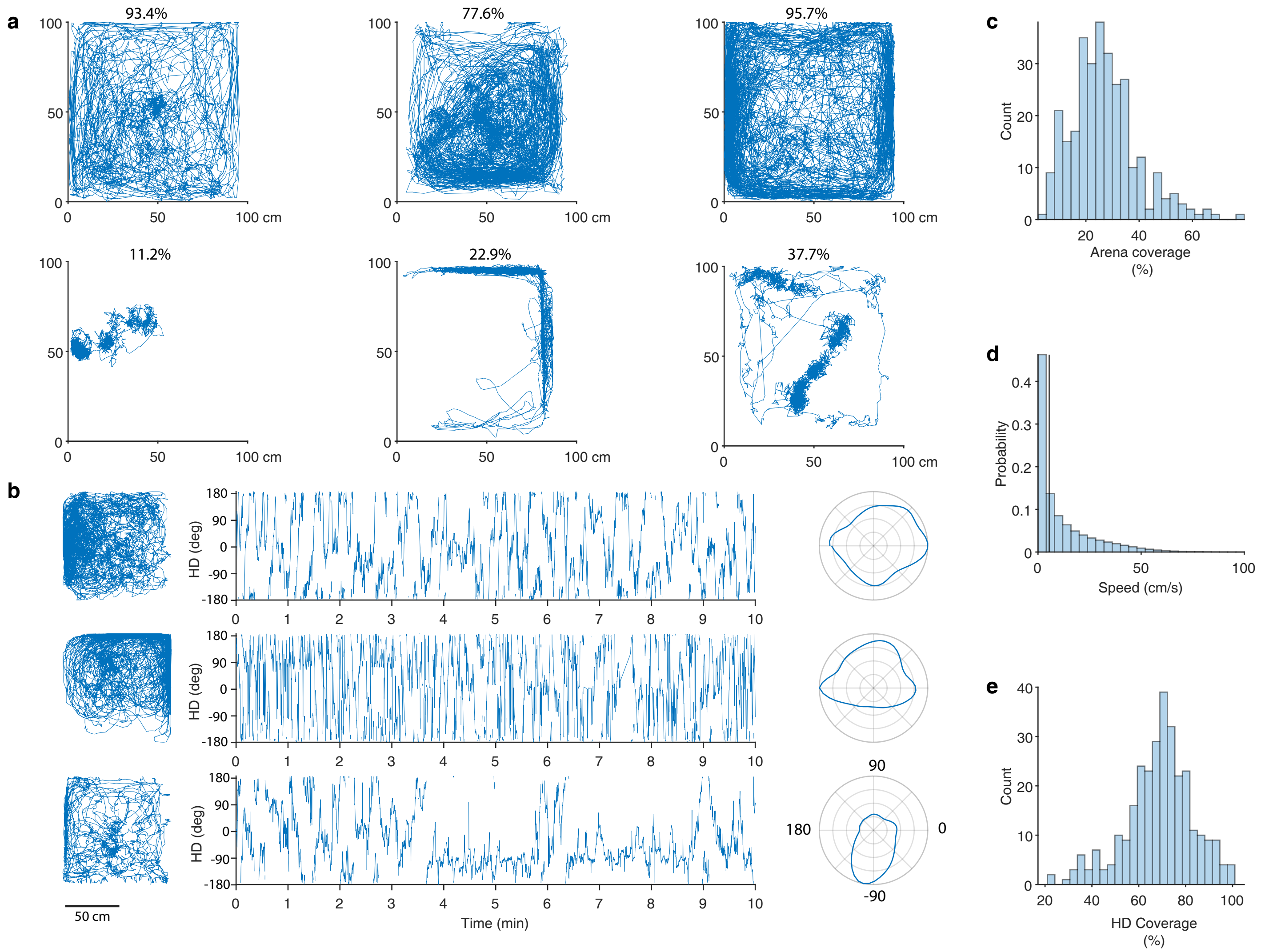


**Fig. S1. Behavioral examples, spatial coverage of the arena, and head-direction coverage. a,** Three examples of sessions with good spatial coverage (top) and three examples of sessions with bad spatial coverage (bottom). The % coverage is indicated above each example. **b,** Head-direction as a function of time from three sessions. Insets on the left show the corresponding position trajectories. Insets on the right show the corresponding polar plots of the behavioral time spent at each head direction. **c,** Spatial coverage of the arena for all behavioral sessions. Percent coverage was computed as described in the Methods. **d,** Probability distribution of the speeds, pooled over all behavioral sessions. The vertical line shows the value of 5cm/sec which was used to divide the data to slower versus faster speeds (Fig. 3h). **e,** The head-direction coverage for all behavioral sessions. Percent coverage is calculated as described in Methods.

Supplementary figure 2

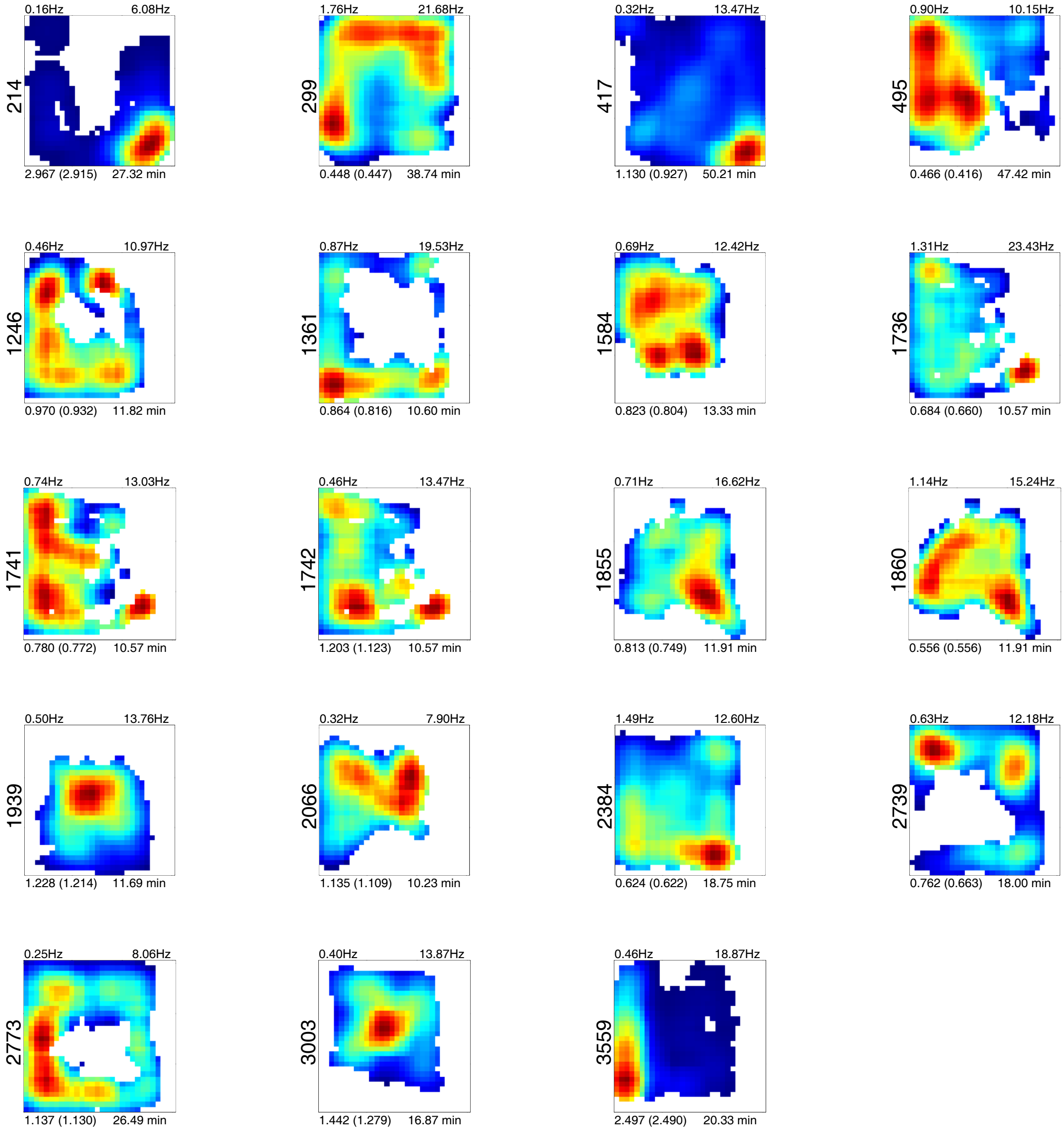


**Fig. S2. Firing-rate maps of 19 cells which showed significant spatial information.** Each square represents the normalized firing-rate map of one cell. Color coded from zero firing-rate (blue) to maximum firing-rate (red). White pixels designate bins that the quail did not visit during the session. Numbers above the plot designate the mean and max firing rates; numbers below the plot designate the spatial information index, in parentheses the information index at the 99th percentile of the shuffled distribution, and on the right is the time duration of the session. Numbers on the left of the panels designate the cell numbers.

### Supplementary figure 3

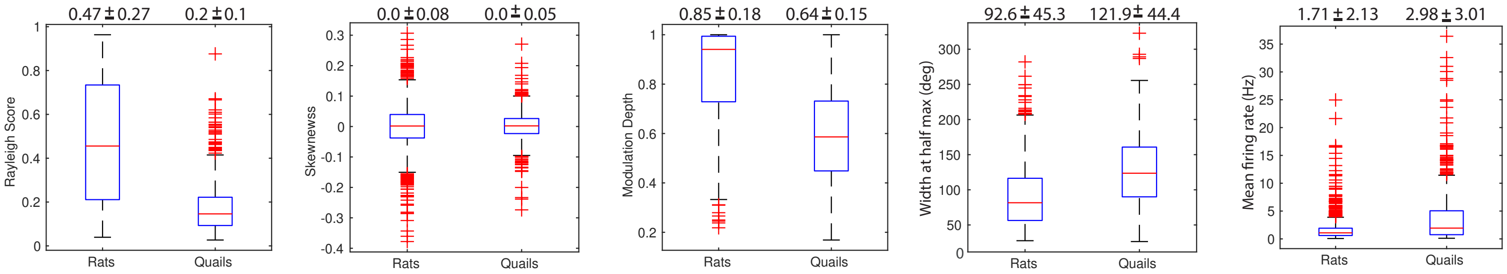


| | Animal | Brain sites | Significant HD cells | Total cells | % HD cells |
|---|---|---|---|---|---|
| Dataset 1* | rats | MEC, PaS, dPrS | 763 | 1300 | 59 |
| Dataset 2** | rats | POR, PaS, dPrS | 283 | 1104 | 26 |
| Dataset 3 | quails | HPF | 260 | 2127 | 12 |

* Data taken from Boccara, et al. Grid cells in pre- and parasubiculum Nat. Neurosci., 13 (2010)

** Data taken from Gofman et al., Dissociation between Postrhinal Cortex and Downstream Parahippocampal Regions in the Representation of Egocentric Boundaries, Current Biology,Volume 29, Issue 16, 2019)

**Fig. S3. Comparison of HD cells recorded in rats with HD cells recorded in quails.** Box plots show the distributions of the Rayleigh vector, skewness. modulation depth, width at half max and mean firing rates in cells recorded from rats compared with cells recorded from quails. Included are cells that pass the criteria for head-direction cells. The same analysis pipeline was used in all cells. MEC- medial entorhinal cortex, PaS - parasubiculum, dPrS - dorsal presubiculum, POR - postrhinal cortex, HPF - hippocampal formation.

Supplementary figure 4

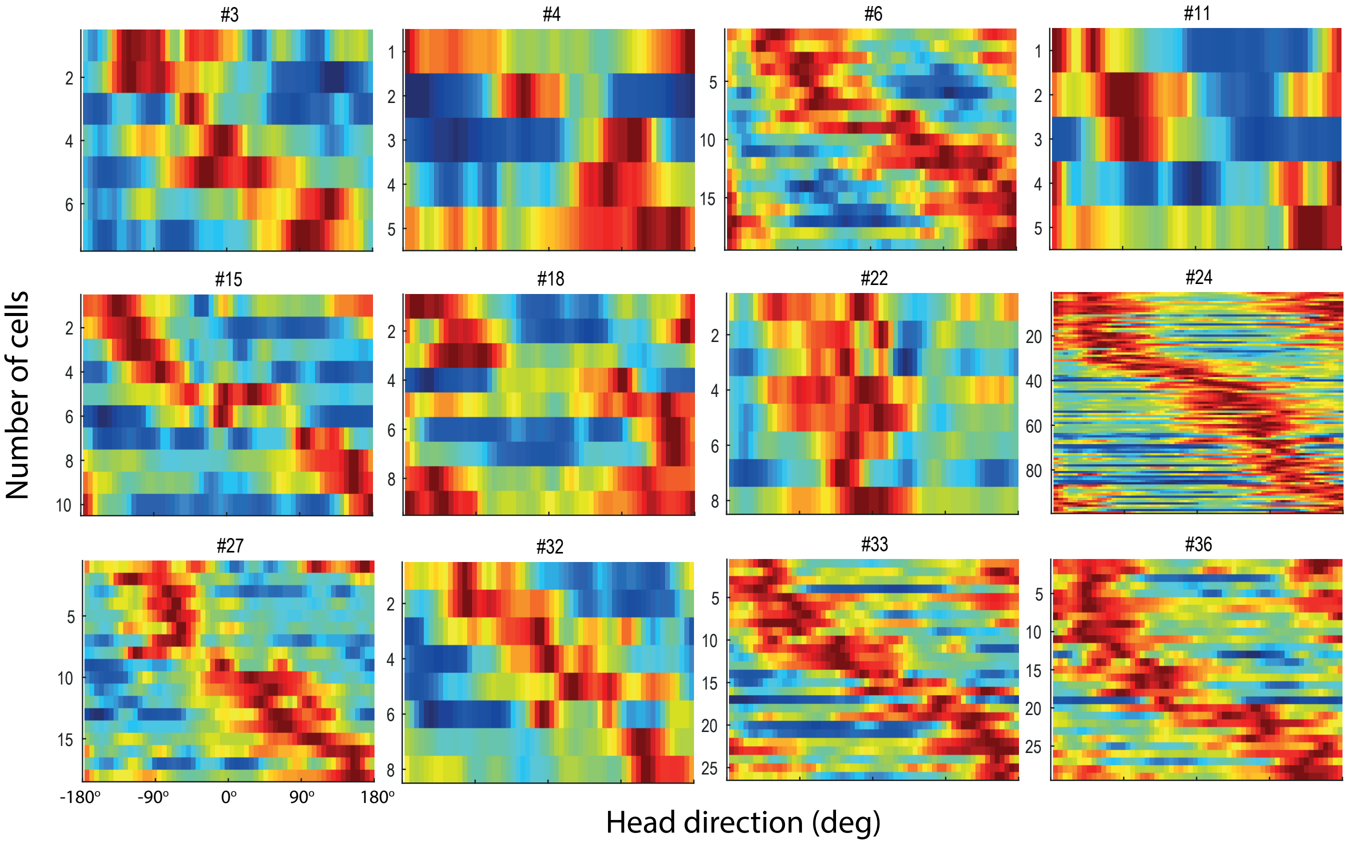


**Fig. S4. Head direction tuning curves from individual quails.** Each plot shows the tuning curves of all significant head-direction cells recorded in one quail; the quail ID number is shown above the plot. Each row designates a single cell and is normalized from zero (blue) to its maximal firing-rate (red). Curves are ordered according to the preferred direction. Shown are data from all the quails in which ≥ 5 significant HD cells were recorded (12 out of 21 quails).

Supplementary figure 5

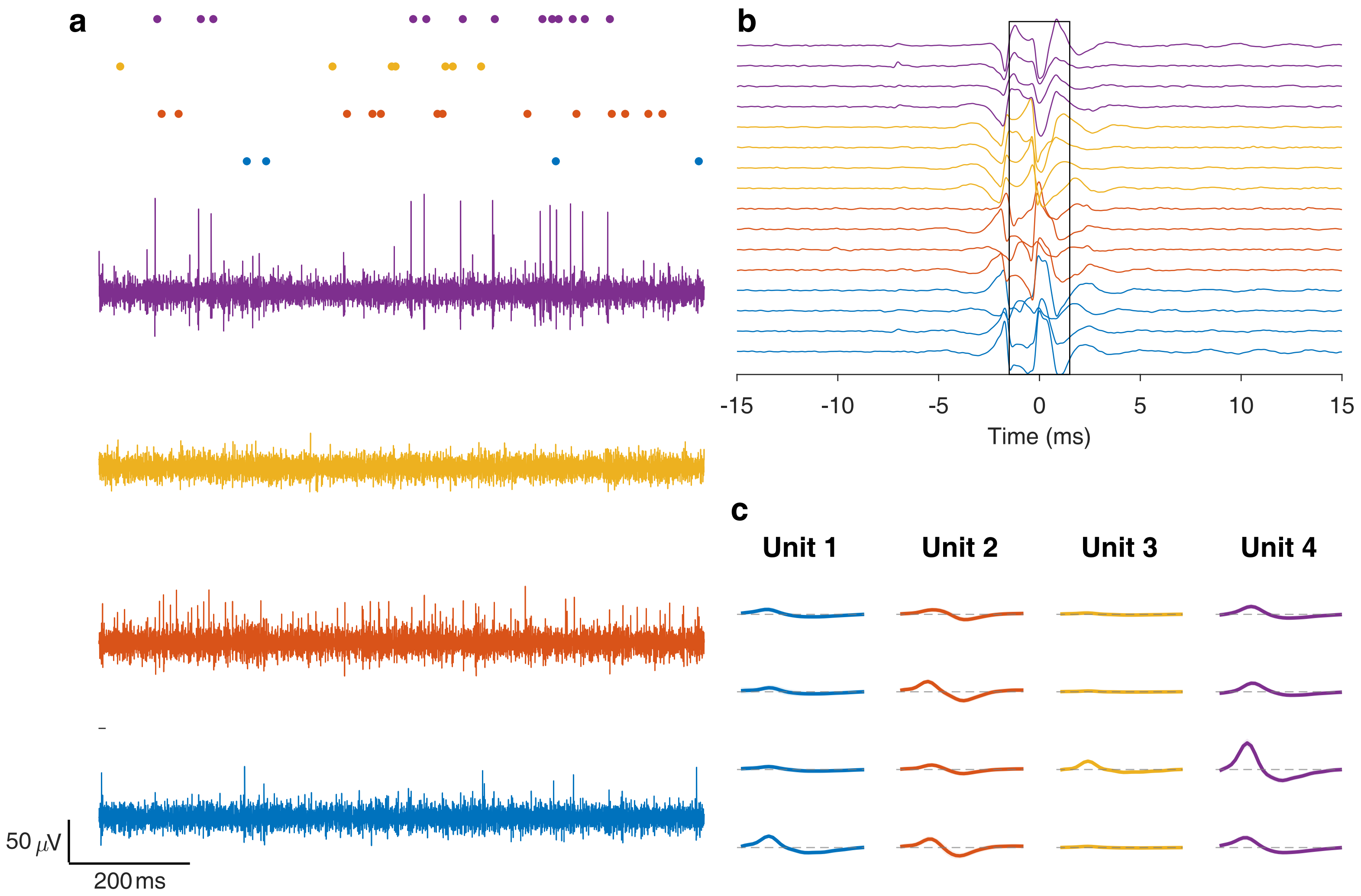


**Fig. S5. Examples of electrophysiological recording traces. a,** Bandpass filtered recording from single electrodes of each tetrode. Top row shows detected spikes, color-coded for each electrode. **b,** Bandpass filtered traces of all 16 channels, showing a movement artifact – usually associated with ground pecking by the bird. Colors designate different tetrodes. Black box marks time window discarded from the recording as part of the artifact-cleaning process. **c,** Spike shapes of the single units shown in panel a. Colors indicate the 4 channels of the tetrode.
